# Spatial Correlation and Breast Cancer Risk

**DOI:** 10.1101/555136

**Authors:** Erin E. E. Fowler, Cassandra Hathaway, Fabryann Tillman, Robert Weinfurtner, Thomas A. Sellers, John Heine

## Abstract

We present a novel method for evaluating the spatial correlation structure in two-dimensional (2D) mammograms and evaluate its merits for risk prediction. Two matched case-control studies were analyzed. Study 1 included women (N = 588 pairs) with mammograms acquired with either Hologic Selenia full field digital mammography (FFDM) units or Hologic Dimensions digital breast tomosynthesis units. Study 2 included women (N =180 pairs) with mammograms acquired with a General Electric Senographe 2000D FFDM unit. Matching variables included age, HRT usage/duration, screening history, and mammography unit. The local autocorrelation function was determined with Fourier analysis and compared with template defined as 2D double-sided exponential function with one spatial extent parameter: n = 4, 12, 24, 50, 74, 100, and 124 defined in pixel widths. The difference between local correlation and template was gauged within a kernel with an adjustable parameter and summarized, producing two measures: the mean (m_n+1_), and standard (s_n+1_). Both adjustable parameters were varied in Study 1. Select measures that produced significant associations with breast cancer were translated to Study 2. Breast cancer associations were evaluated with conditional logistic regression, adjusted for body mass index and ethnicity. Odds ratios (ORs) were estimated as per standard increment with 95% confidence intervals (CIs).

Two measures were selected for breast cancer association analysis in Study 1: m_75_ and s_25_. Both measures revealed significant associations with breast cancer: OR = 1.45 (1.23, 1.66) for m_75_ and OR = 1.30 (1.14, 1.49) for s_25_. When translating to Study 2, these measures also revealed significant associations: OR = 1.49 (1.12, 1.96) for m_75_ and OR = 1.34 (1.06, 1.69) for s_25_.

Novel correlation metrics presented in this work revealed significant associations with breast cancer risk. This approach is general and may have applications beyond mammography.

## 1. Introduction

The spatial correlation properties of images and texture are closely related. For example, a two-dimensional (2D) white noise field is featureless as it lacks spatial correlation. The lack of spatial correlation is a consequence of its flat power spectrum (i.e. all spatial frequency components are statically represented equally). Altering the spectrum of a white noise field by applying a filter produces spatial correlation in the filtered image, perceived as texture. In this work, we present a general method to evaluate the local correlation properties of 2D images based on Fourier analysis. The merits of this approach are evaluated with mammograms, where a specific problem in breast cancer risk is addressed as a possible application. Breast density is a strong breast cancer risk factor, typically estimated from 2D mammograms [1–6]. There are various methods of estimating breast density [3, 6–14]. Most often the focus of these measures is the degree of dense tissue within the breast area. Texture measures also correlate with breast cancer [9]. Although investigated for many years, the connections between breast density and the underlying biological processes are not well understood [15]. Identifying other metrics with defined spatial scales related to breast cancer could be useful for informing future studies designed to understand the related biological processes with breast structure, in addition to risk prediction purposes.

## 2. Materials and Methods

### 2.1 Population and Imaging

This study includes two populations derived from the same geographical region over distinct time frames. For the purposes of this report, we refer these as Study 1 and Study 2 to align with the order of our current presentation and data processing sequence (i.e. not related to the study timeframes). Study 1 included 588 individually matched case-control pairs of women that attended the breast clinics at Moffitt Cancer Center (MCC) between 2013 and 2018. Study 2 included 180 individually matched case-control pairs that attended the breast clinics at MCC between 2007 and 2011 presented previously [16]. For Study 1, 2D study mammograms were acquired from one of six Hologic (Hologic, Inc., Bedford, MA) mammography units: three conventional 2D Selenia full-field digital mammography (FFDM) units and three Dimensions digital breast tomosynthesis (DBT) units. These units use direct x-ray detection and have 70μm pitch. For Study 2, 2D study mammograms were acquired with one General Electric (General Electric Medical Systems, Milwaukee, WI, USA) Senographe 2000D FFDM unit. This unit has 100μm pitch. In both studies, raw data was used for analytical purposes; these images are in monochrome 1 format (i.e. dense tissue has lower intensity values than adipose tissue) with 14-bit dynamic range per pixel. For illustration purposes, we used for presentation (processed) images when noted; these are the images used for clinical purposes. The unaffected breast was used as the study image for cases. A given control study image laterality was determined by matching with its respective case’s study image side. Processing was constrained to the largest rectangular box that can be inscribed within the breast area as illustrated in Figure 1 using an automated algorithm developed and described in detail previously [16, 17].

**Figure 1:**
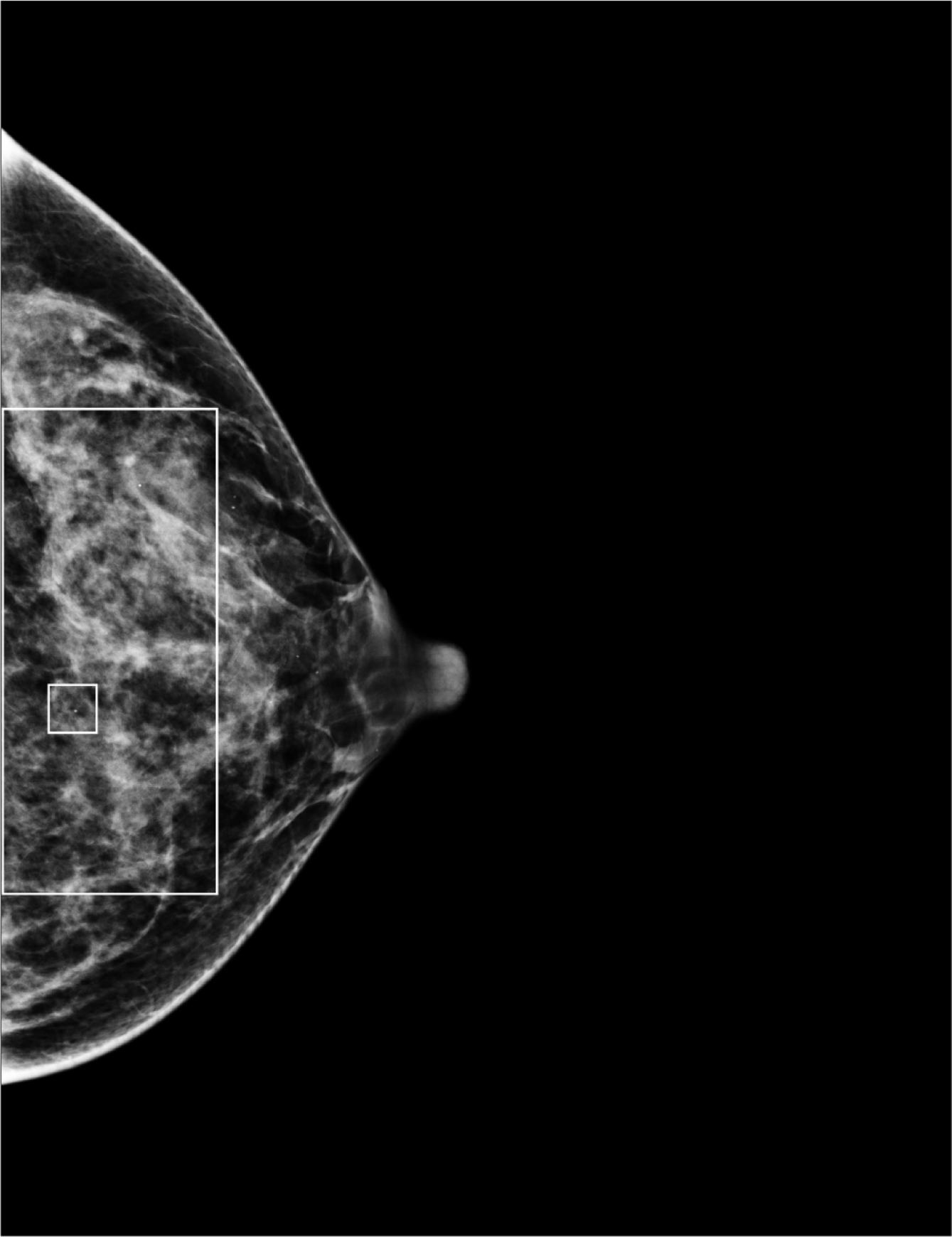
Mammogram with the largest rectangle: This shows typical mammogram in processed format used for viewing purposes with the largest rectangle inscribed within the breast area. The smaller region is 125×125 pixels equating with n = 124. We used this smaller region for illustration purposes in Figure 1 and 2.

Both studies employed the same protocol. Cases had a first-time diagnosis, pathology verified, of unilateral breast cancer. These cases were either (i) women diagnosed with breast cancer attending the breast clinics at MCC or (ii) attendees of surrounding area clinics sent to MCC for breast cancer treatment or diagnostic purposes and found to have breast cancer. Controls were attendees of MCC without a history of breast cancer. Controls were individually matched to cases on age (±2 years), hormone replacement therapy (HRT) usage and current duration, screening history, and mammography unit. The HRT match was based on status of current users or non-users. Non-users included women that have not taken HRT for at least two years. If a case was a current HRT user, the control was matched on this duration (±2 years). Controls were matched by screening history using a three-category classification. Group 1 included women with prior screening history by any means; the duration between the last screening and the study image date must be no more than 30 months. Group 2 included women without screening history. Group 3 included women with a screening history that does not fit within in Group 1 or Group 2. We used mammograms in cranial caudal (CC) orientation as study images. The unaffected breast was used as the study image for cases (image acquired before treatment) and the matching lateral breast for controls.

### 2.2 Image Measures

This new approach was based on comparing the local autocorrelation properties of a given mammogram with a template and then summarizing the local measures within a given image. For reasons illustrated below, we defined the correlation template as a 2D double sided symmetric exponential function expressed as 
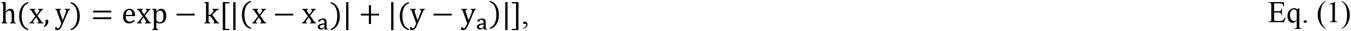
where x and y are pixel coordinates (i.e. integers) in a Cartesian coordinate system, and k is the decay constant. We used h(x, y) to make comparisons with local autocorrelation function in mammograms about an arbitrary location expressed as (x_a_, y_a_). The local image region centered about this location is defined as f(x, y) with x and y ranging from (x_a_ - n, x_a_ + n) and (y_a_ - n, y_a_ + n), respectively, giving a region size of (2 × n + 1) × (2 × n + 1) pixels, where n is an even integer. In practice, n was an experimental parameter to be determined. The associated empirical local autocorrelation function, defined as h_e_(x, y), was determined using Fourier transform (FT) relationships by invoking the 2D autocorrelation theorem [18] and was forced to have the same spatial extent in each coordinate direction ranging from –n through n (demonstrated below). The parameter k is a function of n, derived empirically to ensure that h(x, y) tends to zero expressed as 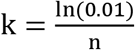

Modification to the region dimensions was required to eliminate wraparound effects of the discrete FT and force the associated relationships to match those derived from the continuous FT theory. This modification is illustrated with a specific example. Figure 2 (left) shows an image region with n = 124, corresponding to the smaller region shown in Figure 1. We extended the spatial extent of this region by zero padding by 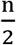 i.e. 62) about f(x, y) in each spatial direction as illustrated in Figure 2 (left) and then used the 2D Fourier correlation relationship [18] to determine h_e_(x, y). We take the FT of f(x, y) after zero padding, giving F(f_x_, f_y_). We then formed the power spectrum expressed as F(f_x_, f_y_)×F^*^(f_x_, f_y_); the asterisk indicates complex conjugate and f_x_, and f_y_ are Cartesian spatial frequency coordinates in x and y directions, respectively. This was followed by Fourier inversion giving the empirical autocorrelation function h_e_(x, y) illustrated in Figure 2 (middle). This empirical correlation function was then normalized such that h_e_(x_a_, y_a_) = 1. The multiplication in the Fourier domain used to form the power spectrum is equivalent to performing a correlation operation in the image domain[18]. When viewed in the image domain, zero padding ensures that the shifted function in the correlation operation moves into a region where the stationary (non-shifted) function is zero. This accounts for the wraparound artifact in discrete FT, allowing a closer relationship to continuous FT theory. This is illustrated Figure 2 (right), which shows a profile through h_e_(x, y_a_). Note that the correlation tends to zero and the spatial extent along the profile spans from shift = −124 to shift = 124. In contrast, Figure 3 shows the same example region without zero padding (left), the related h_e_(x, y) (middle), and profile through h_e_(x, y_a_) [right]. Note that the profile does not tend to zero and that the spatial extent spans from [−64, 64] measured in pixel width shifts.

**Figure 2:**
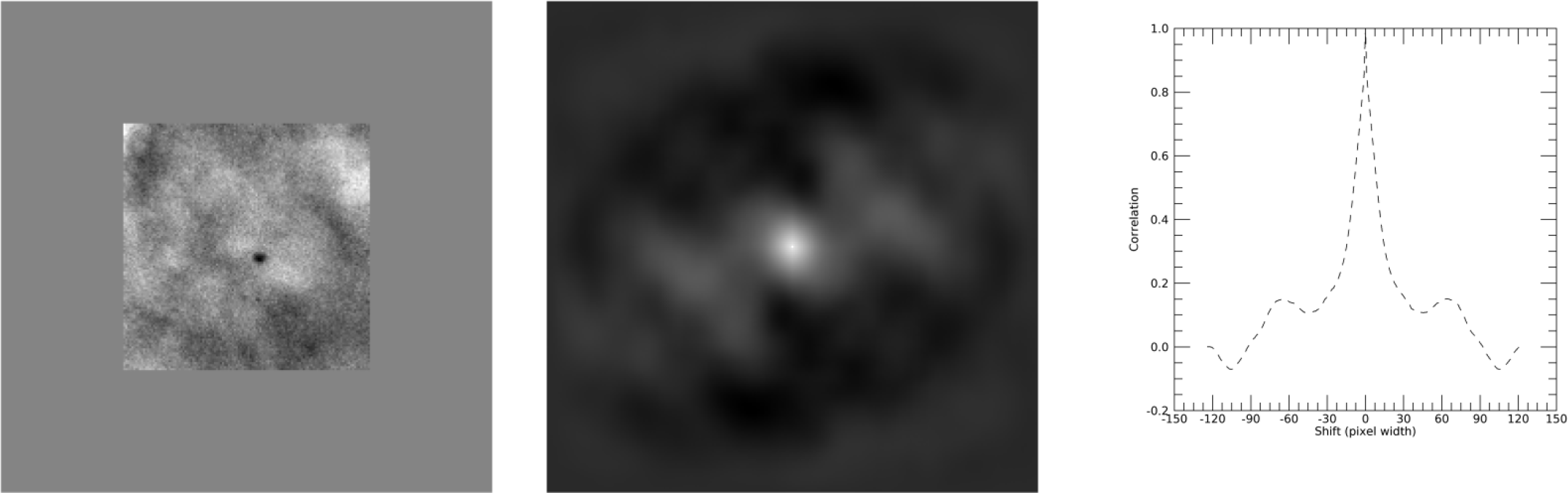
Correlation with zero padding: The section on the left was extracted from Figure 1 and zero padded. In this example, n = 125, giving a region of (2n+1)×(2n+1)= 249×249 pixels with the zero padding. The middle pane shows the corresponding two-dimensional auto correlation function, h_e_(x, y). The pane on the right shows a slice along the middle of h_e_(x,y) along the x-direction (horizontal). Note that h_e_(x, y) tappers to zero as the shift approaches |124|.

**Figure 3:**
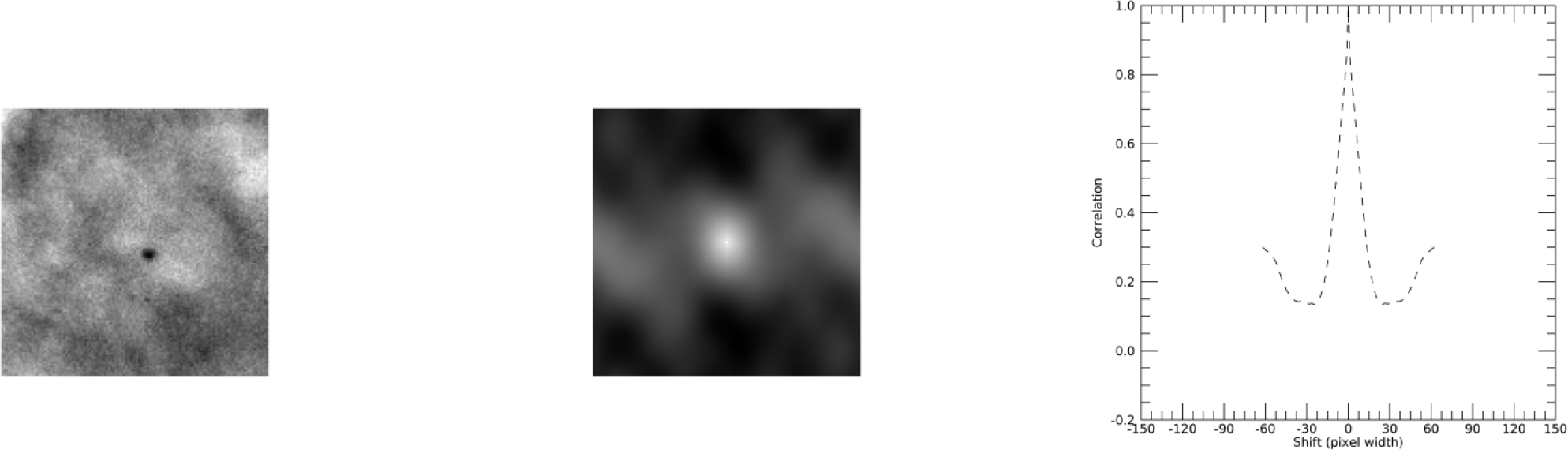
Correlation without zero padding: The section on the left was extracted from Figure 1. In this example, n = 125, giving a region of (n + 1) × (n + 1) = 125×125 pixels. The middle pane shows the corresponding two-dimensional autocorrelation function, h_e_(x, y). The pane on the right shows a slice along the middle of h_e_(x, y) along the x-direction (horizontal). Note that h_e_(x, y) does not tapper to zero in contrast with the example shown in Figure 2.

The local autocorrelation function, h_e_(x,y), was estimated as a function of location across each mammogram and compared with the template using a kernel function expressed as 
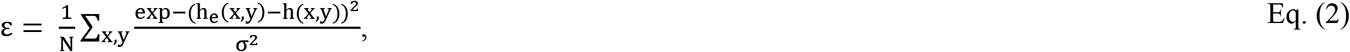
resulting in a spatial map of the local correlation. The comparison within the kernel normalizes ε by constraining it between (0, 1). Both the mean and standard deviation (SD) of the ε distribution for a given image was used as two measures defined as ε_m_ and ε_s_ respectively. For a given n, specific ε_m_ and ε_s_ metrics were labeled as m_n+1_ and s_n+1_, respectively. The kernel width, σ, is the other experimental parameter to be determined. To reduce processing time, the coordinates (x_a_, y_a_) were shifted across the image in 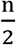 steps in both spatial dimensions.

The operator assisted software, Cumulus 3, was implemented by JH to determine the percentage of breast density (PD) for processed images for both studies. The PD findings for Study 2 were discussed previously [16] and are presented in this report as well for completeness. Cases and controls were randomly mixed and JH was blinded to all patient information. PD was used as a standard for comparison with selected correlation metrics.

### 2.3 Statistical Methods

Study 1 was used for exploration purposes. Seven region sizes for n+1 were investigated (expressed in pixel widths): 5, 13, 25, 51, 75, 101, and 125, corresponding with approximately 0.35mm, 0.91mm, 1.75mm, 3.57mm, 5.25mm, 7.07mm, and 8.75mm, respectively. For each region size, σ was varied from 0.001 through 0.05 with 0.001 increments giving 50 separate settings. To demonstrate that the template is a reasonable choice, a random selection experiment was used for illustration. Five mammograms were selected at random from the control group in Study 1, and the largest rectangle was determined for each mammogram. For each of these regions, one position was selected at random and h_e_(x, y) was constructed for n + 1 = 51. For each random h_e_(x,y) graphical comparisons were made with the respective h(x, y) by plotting profiles taken through both functions through the coordinate axes.

The spatial correlation processing will result in many measures per image. There are 7 region sizes each with 50 settings for σ and two measurements giving a total of 7 × 50 × 2 = 700 spatial correlation measures for each mammogram. To acquire an overview of these measures, we applied a paired t-test across case and control groups for each measure and summarized these measures according to three significance levels: 0.05, 0.025, and 0.01. A two-dimensional correlation table was developed for select measure(s) with p < 0.01. Due to the way the spatial correlation measures were developed, we expect high correlation between them. Measures were selected for further analysis based on this significance level, correlation (Pearson’s correlation coefficient denoted by R) between the other template measures and PD: maximal difference between case and control groups, minimal correlation with PD, and minimal correlation with other correlation measures. These select measure(s) were analyzed with conditional logistic regression. The specific settings (n and σ) for these select measure(s) were used to process Study 2 mammograms. To keep the spatial distances the same, the region size(s) determined with Study 1were translated to Study 2 by reducing the respective region dimension by the pitch ratio,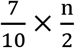, with rounding.

Conditional logistic regression modeling was used to estimate breast cancer associations. Image measurement distributions were log-transformed. Odds ratios (ORs) were used as the association metrics with 95% confidence intervals (CIs). Continuous ORs were estimated as per standard deviation (SD) increment determined for the respective image measurement distribution. We also considered the area under the receiver operating characteristic curve (Az) for each model with 95% CIs. Models will be presented unadjusted and adjusted for body mass index (BMI) and ethnicity. In Study 1, the case/control paired t-test for BMI was calculated with the 581 complete pairs due to missing data on seven controls: BMI values were set to the control distribution BMI mean for modeling purposes. When comparing proportions, McNemar’s (exact) test was used for within population comparisons. When comparing continuous measures, we used the t-test. Image processing was implemented with IDL version 8.6 (Exelis Visual Information Solutions, Boulder, CO) and regression analyses was done using SAS version 9.4 (SAS Institute Inc., Cary, NC).

## 3. Results

Patient characteristics for Study 1 and Study 2 [16] are provided in Table 1a and 1b, respectively. Both studies were comprised of primarily Caucasian women (83.8% for Study 1 and 89.2% for Study 2). In both studies ethnicity was similar with Non-Hispanics representing 84.3% in Study 1 and 90.6% in Study 2. The Hispanic population was higher in controls than cases in Study 1 (p < 0.0001) and similar in Study 2. The mean age of participants in either study was approximately 58 years. Cases had a higher BMI than controls with p = 0.0003 and p = 0.0071 for Study 1 and Study 2, respectively. Menopausal status was similar between cases and controls with p = 0.50 for Study 1 and p = 0.076 for Study 2 when comparing menopausal women versus other status (i.e. pre-menopausal plus unknown). PD was similar between cases and controls in Study 1 (p = 0.25) and significantly different in Study 2 (p = 0.025).

**Table1a:**
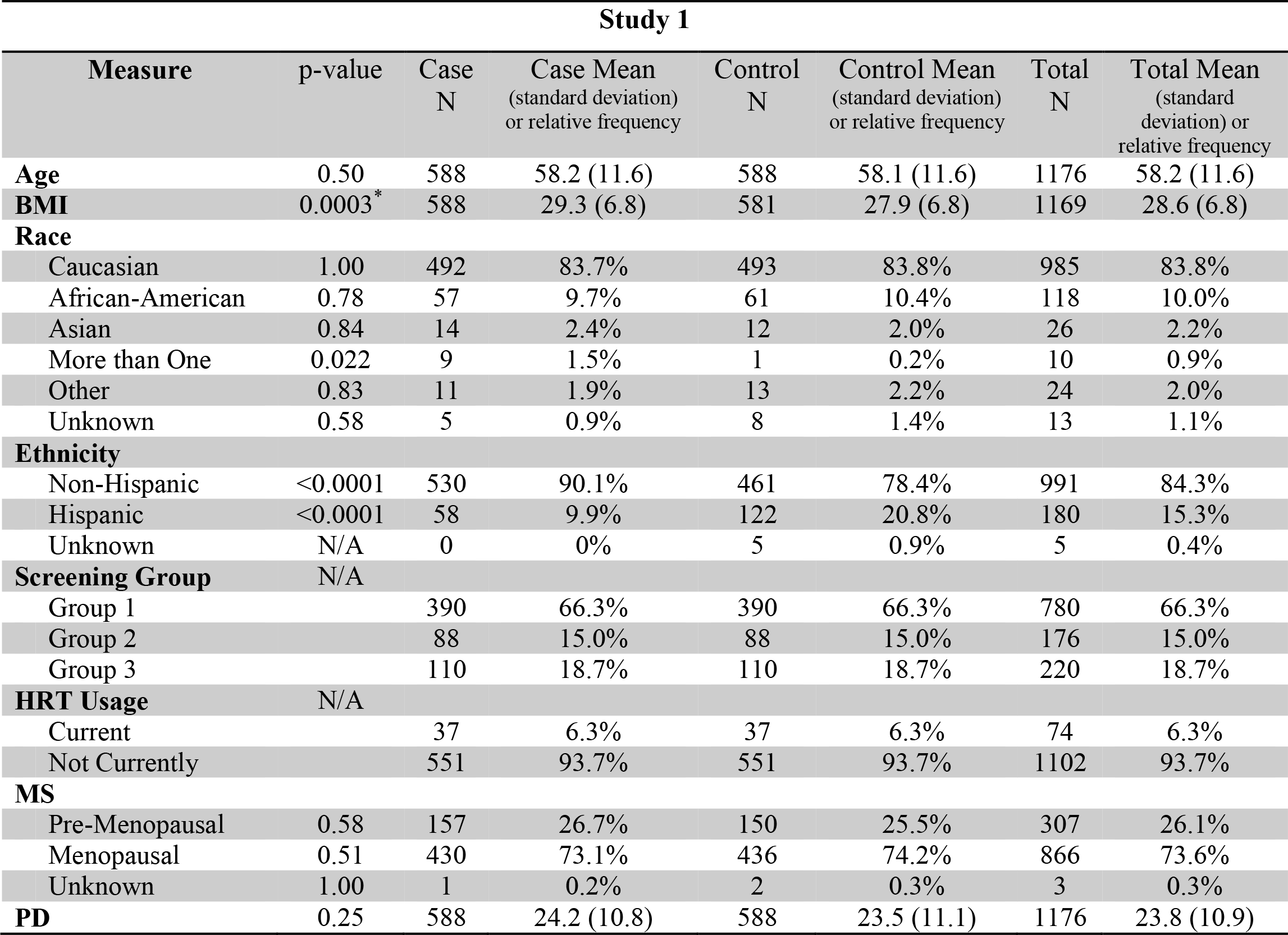
Study 1 characteristics: This table provides Study 1 characteristics by either distribution mean for a given measure or percentages of the population. Where applicable, the standard deviation of the respective distribution is provided parenthetically. Images were acquired with either Selenia FFDM or Dimensions DBT units. The case/control paired t-test for BMI was calculated with the 581 complete pairs due to missing data on seven controls.

**Table1b:** Study 2 characteristics. This table provides Study 2 population characteristics by either distribution mean for a given measure or percentages of the population. Where applicable, the standard deviation of the respective distribution is provided parenthetically. Images were acquired with a Senographe 2000D unit.

Study 2
| Measure | p-value | Case<br>N | Case Mean<br>(standard deviation)<br>or relative frequency | Control<br>N | Control Mean<br>(standard deviation)<br>or relative frequency | Total<br>N | Total Mean<br>(standard deviation)<br>or relative frequency |
| --- | --- | --- | --- | --- | --- | --- | --- |
| **Age** | 0.25 | 180 | 58.6 (10.5) | 180 | 58.5 (10.4) | 360 | 58.6 (10.4) |
| **BMI** | 0.0071 | 180 | 26.6 (4.7) | 180 | 25.2 (4.3) | 360 | 25.9 (4.5) |
| **Race** |  |  |  |  |  |  |  |
| Caucasian | 0.74 | 159 | 88.3% | 162 | 90.0% | 321 | 89.2% |
| African-American | 0.26 | 7 | 3.9% | 13 | 7.2% | 20 | 5.6% |
| Asian | 0.291 | 7 | 3.9% | 3 | 1.7% | 10 | 2.8% |
| More than One | N/A | 2 | 1.1% | 0 | 0.0% | 2 | 0.6% |
| Other | N/A | 4 | 2.2% | 0 | 0.0% | 4 | 1.1% |
| Unknown | 1.00 | 1 | 0.6% | 2 | 1.1% | 3 | 0.8% |
| **Ethnicity** |  |  |  |  |  |  |  |
| Non-Hispanic | 0.85 | 164 | 91.1% | 162 | 90.0% | 326 | 90.6% |
| Hispanic | 0.84 | 14 | 7.8% | 16 | 8.9% | 30 | 8.3% |
| Unknown | 1.00 | 2 | 1.1% | 2 | 1.1% | 4 | 1.1% |
| **Screening Group** | N/A |  |  |  |  |  |  |
| Group 1 |  | 162 | 90.0% | 162 | 90.0% | 324 | 90.0% |
| Group 2 |  | 5 | 2.8% | 5 | 2.8% | 10 | 2.8% |
| Group 3 |  | 13 | 7.2% | 13 | 7.2% | 26 | 7.2% |
| **HRT Usage** | N/A |  |  |  |  |  |  |
| Current |  | 36 | 20.0% | 36 | 20.0% | 72 | 20.0% |
| Not Currently |  | 144 | 80.0% | 144 | 80.0% | 288 | 80.0% |
| **MS** | 0.076 |  |  |  |  |  |  |
| Pre-Menopausal |  | 38 | 21.1% | 48 | 26.7% | 86 | 23.9% |
| Menopausal |  | 142 | 78.9% | 132 | 73.3% | 274 | 76.1% |
| **PD** | 0.025 | 180 | 22.6 (14.7) | 180 | 19.8 (12.9) | 360 | 21.2 (13.9) |

To justify the template choice, the random selection illustrations are discussed first. The top row of Figure 4 shows the largest rectangles for five mammograms that were selected at random (processed images are used for display purposes only). The smaller outlined box (n+1 = 51) within each image was also selected at random. Figure 5 shows five randomly selected h_e_(x, y) images for each smaller region with the same ordering as Figure 4. The middle row of Figure 5 shows profiles through the origin along the x-direction and the bottom row shows profiles along the y-direction for h(x,y) [solid line] and h_e_(x, y) [dashed line]. Comparing the empirical profiles with the corresponding template profiles illustrates the similarities graphically. The bottom row of Figure 4 shows the related spatial maps of the local correlation difference measure, ε, derived from Eq. (3). Brighter regions represent stronger agreement with the reference template.

**Figure 4:**
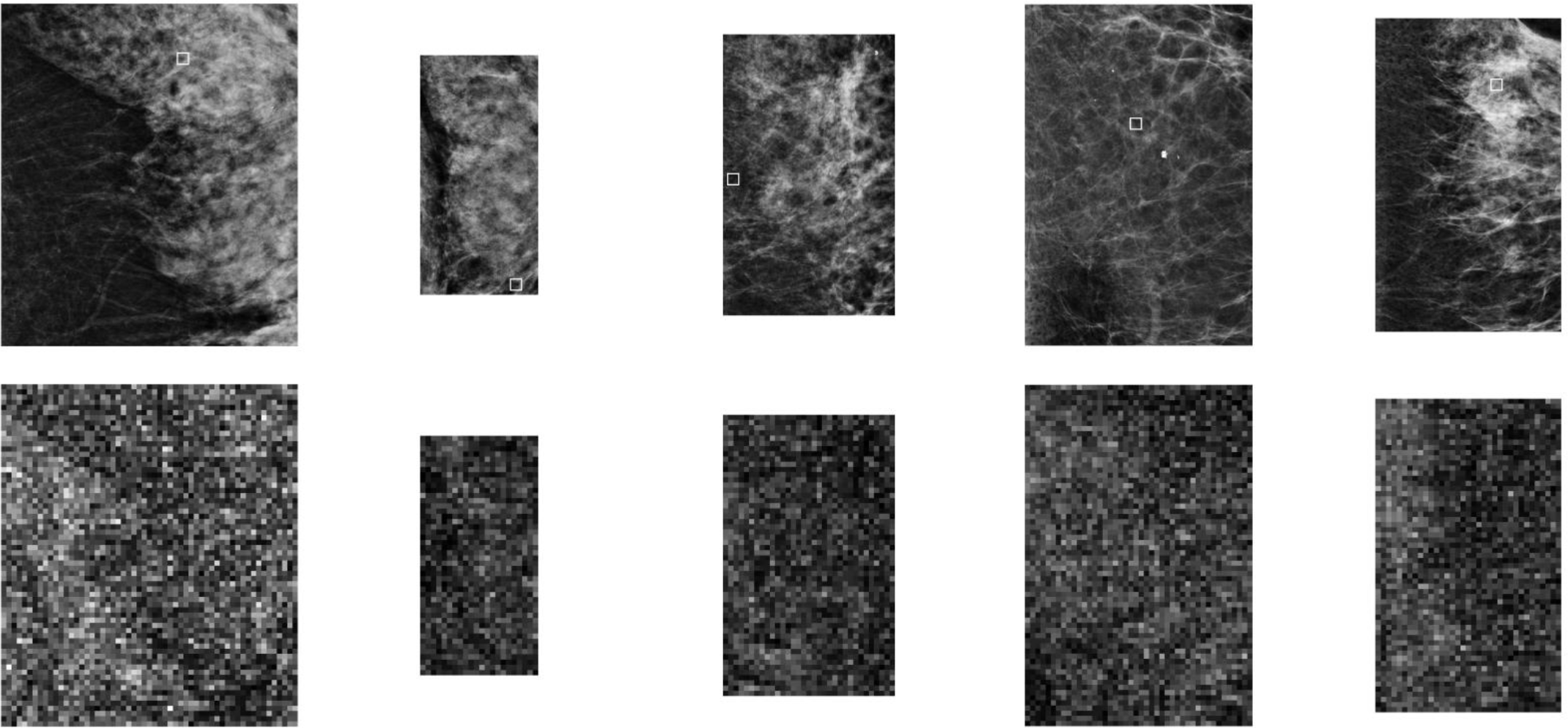
Random mammogram section illustrations and spatial autocorrelation maps. This figure shows the largest rectangles constructed from five randomly selected mammograms (top row). The dimensions in pixels (left to right) are: 1470×1698, 586×1187, 852×1394, 1128×1693, and 921×1555. The small box (n+1 = 51) outlined within each image was also selected at random. The respective spatial distributions (resized to the full rectangle dimensions) for ε are illustrated in the bottom row; both the far left and far right images illustrate clearly that adipose regions (dark areas in the top row images) correspond with larger ε (bright areas in the bottom row images).

**Figure 5:**
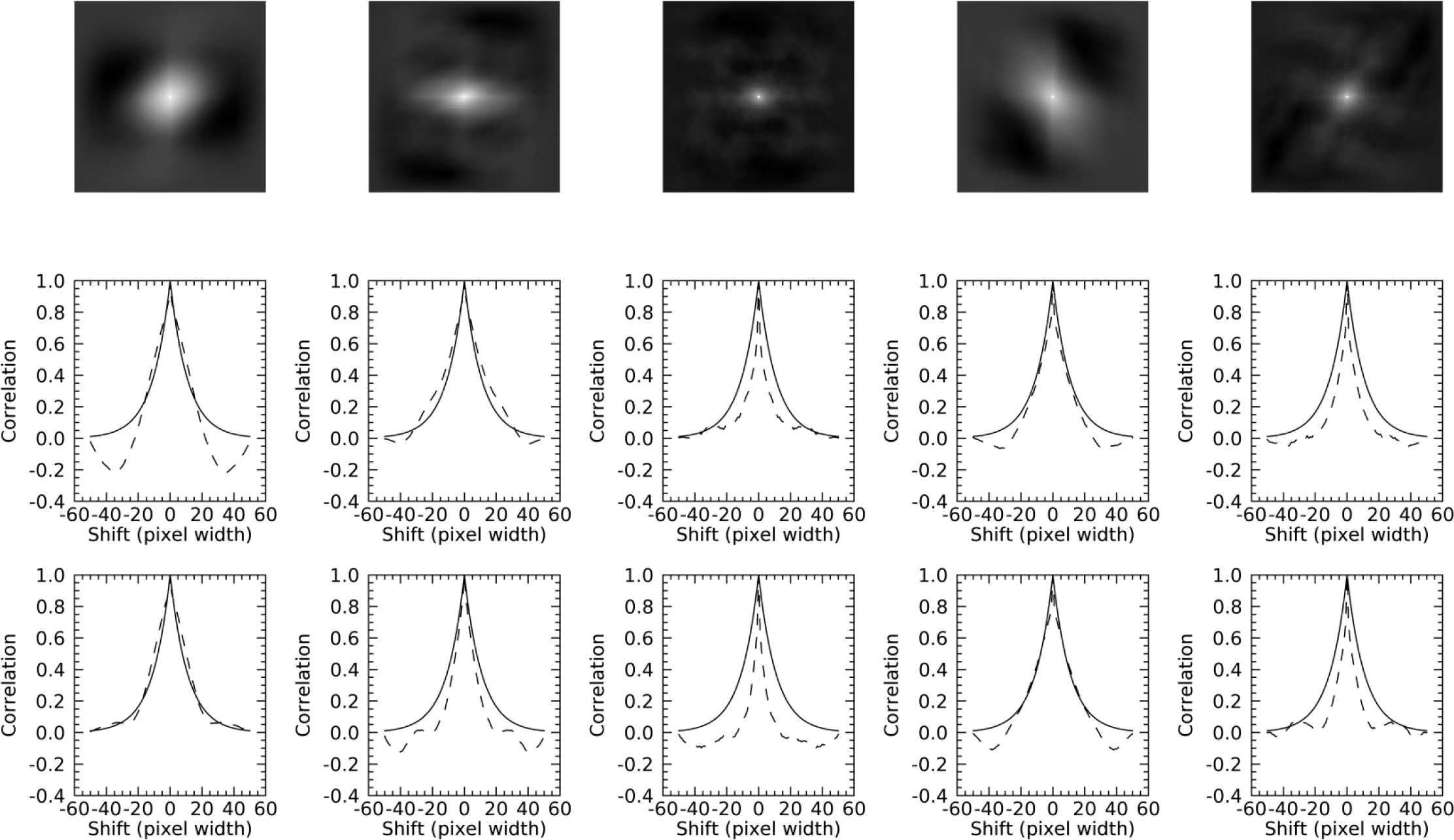
Randomly selected autocorrelation images and profiles. The top row shows the associated h_e_(x,y) images for the randomly selected regions (n + 1 = 51) shown in Figure 4 (smaller region in the top row). The middle row shows the respective profiles taken through h_e_(x, y) [dashes] and h(x,y) [solid] along the x axis. The bottom row shows the respective profiles taken through the y axis.

Table 2 shows the initial evaluation derived from the t-test. These findings are separated by measurement-type, significance levels, and by distribution stochastic dominance. Most of the mean-measurements were significant at both p < 0.05 and p < 0.025. When considering p < 0.01, both m_51_ and m_75_ were significant for all σ settings as was s_13_ indicating these measures were robust with respect to this parameter setting. Table 3 shows the correlation measures with p < 0.01 for the σ setting that gave the smallest p-value. There was a high degree of correlation across the m_n+1_ measures with R > 0.95 for all pairs, whereas the correlation between the s_n+1_ measures varied with R ~ 0.80-0.86 for most pairs; deviation was noted between s_5_ and s_25_ with R = 0.53. The correlation across the m_n+1_ and s_n+1_ measures was negative and varied with R ~ (−0.86 - −0.63). As such, we choose the optimal m_75_ (5.75mm) determined with σ = 0.006 (p = 0.005) as one measure to examine more closely and translate to Study 2. The optimal sigma setting for s_25_ was determined with σ = 0.050 (p < 0.002), where the case distribution exhibited stochastic dominance. The correlation between m_75_ and s_25_ was R = −0.68; the respective correlation between these measures with PD was R = −0.70 and R = 0.50, respectively. As such, we also selected s_25_ (1.75mm) to examine in more detail as another measure.

**Table 2:**
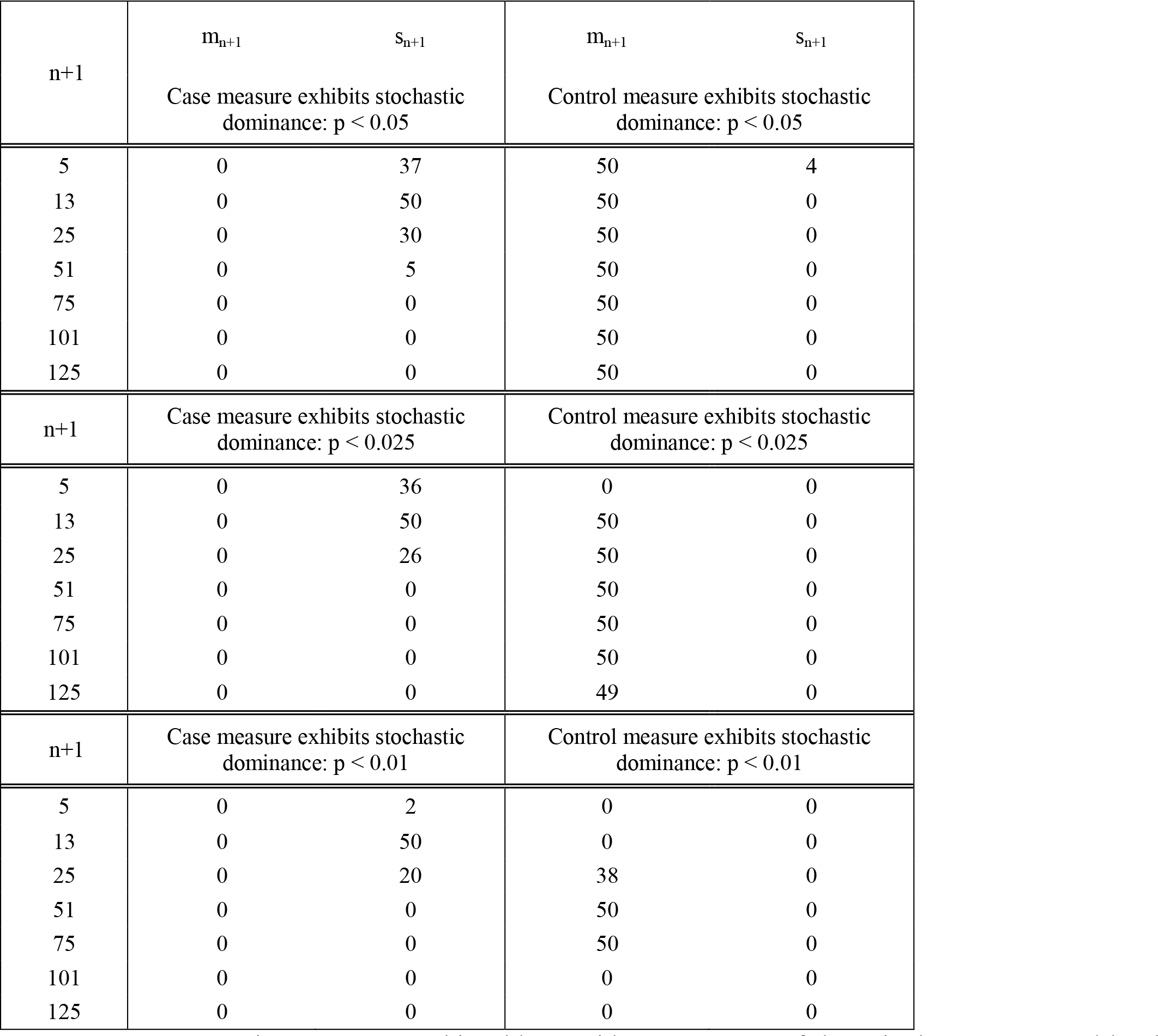
t-test comparison summary. This table provides a summary of the paired t-test separated by the two metrics, case or control distribution stochastic dominance, and three significance levels. The spatial correlation dimensional parameter, n+1, is provided in the left column.

**Table 3:**
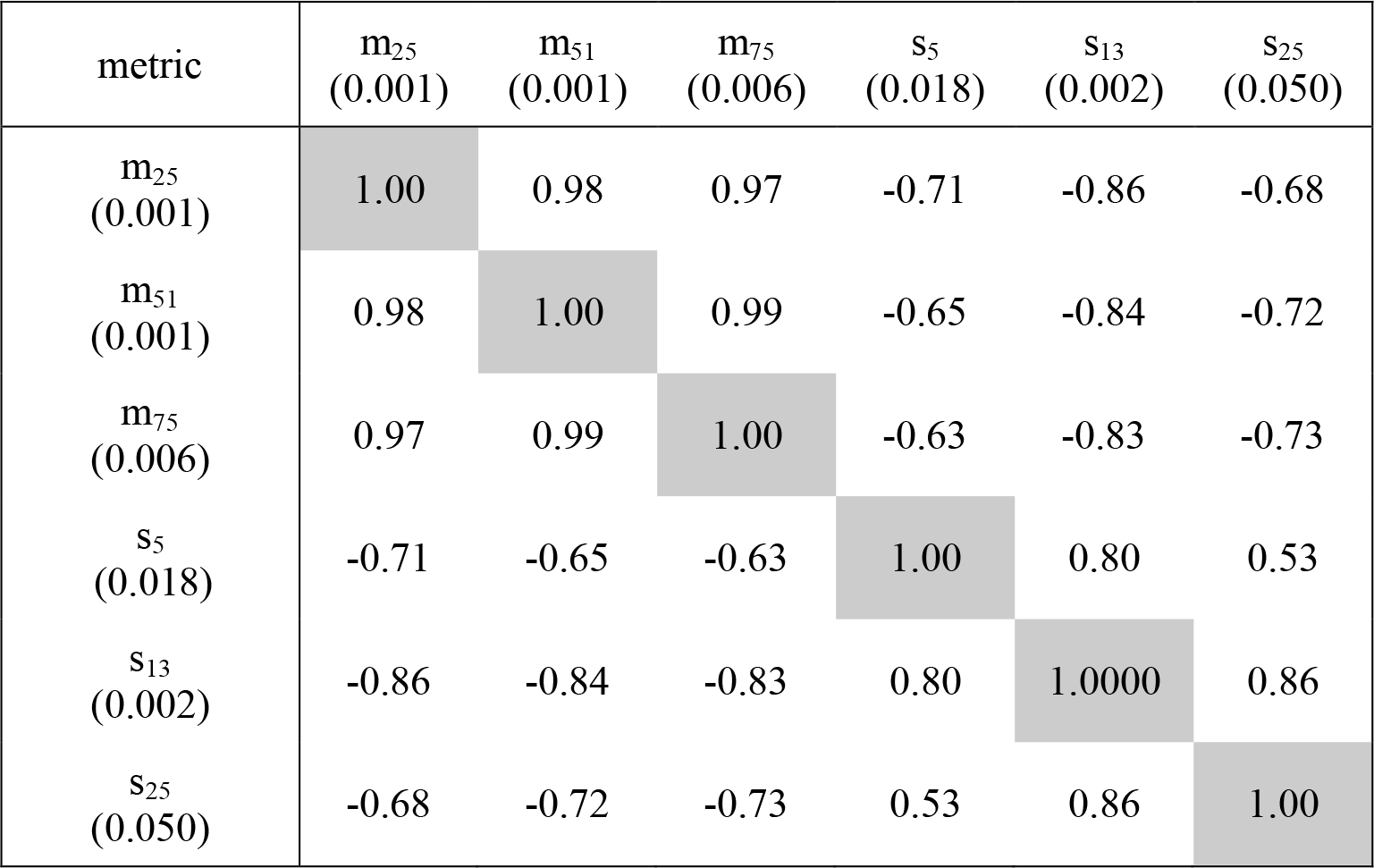
Correlation table. This table provides a summary of the linear correlation coefficients for measures in Table 2 with p < 0.01. The σ settings are provided parenthetically for the minimal p-value.

Table 4 (top) shows the breast cancer associations provided by PD (left), m_75_ (middle) and s_25_ (right) for Study 1. All measures provided significant association with breast cancer in the adjusted models. Cases exhibited stochastic dominance for both PD and s_25_ and both methods produced similar ORs of 1.34 and 1.30, respectively with Az = 0.64 and 0.63. In contrast controls exhibited stochastic dominance for m_75_ with OR = 0.69 with Az = 0.64, which was similar to the Az produced by the other measures. Inverting this measure gave OR = 1.45 (1.23, 1.66) showing magnitude similarity with other associations. We translated m_75_ and s_25_ to Study 2 giving n+1 = 53 and n+1 = 17 respectively, preserving spatial distances and metrics across studies. The corresponding Study 2 breast cancer associations for PD, m_75_ (m_53_) and s_25_ (s_17_) are provided in the Table 4 (bottom). All measures provided significant associations in the adjusted models. Cases exhibited stochastic dominance for both PD and s_25_ with ORs of 1.73 and 1.34, respectively with Azs of 0.64 and 0.63. When comparing respective metrics across Studies (adjusted models), PD provided a marginally greater OR in Study 2 with an identical Az. The remaining metric, m_75_, had an OR 0.67 with an Az of 0.61. Inverting the associations for m_75_, gave OR = 1.49 (1.12, 1.96). Both m_75_ and s_25_ provided nearly identical ORs in both studies, whereas Az was marginally greater for m_75_ in Study 1 (0.64 vs. 0.61) and was identical for s_25_.

**Table 4:**
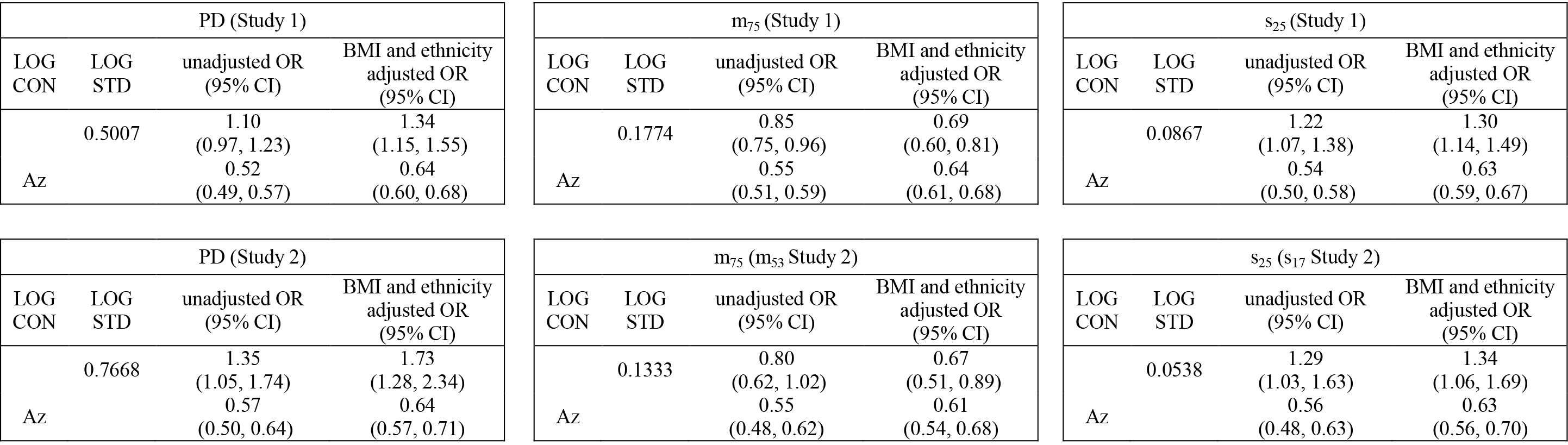
Conditional logistic regression modeling for Study 1 and Study 2. This table provides odd ratios (ORs) and the area under the receiver operating characteristic curve (Az) with 95% confidence intervals (CIs) for PD and the two selected spatial correlation metrics for both studies: m_75_ (or equivalently m_53_ for Study 2) and s_25_ (or equivalently s_17_ for Study 2). Models are provided in unadjusted and adjusted for body mass index (BMI) and ethnicity formats.

## 4. Discussion

A technique for evaluating the correlation structure in mammograms based on a template comparison was presented. This technique was applied to one study for exploration purposes by varying the adjustable parameters. Many measures were significantly different across the case and control groups. Two correlation measures were selected for further analytical scrutiny. These measures produced similar and significant breast cancer associations when applied in both studies; indicating that they are robust given the different FFDM technologies. Moreover, the findings from Study 2 did not require additional adjustments beyond accounting for pitch differences. The first and last images in Figure 4 show that adipose tissue may have a stronger association with the reference than glandular tissue, which is also indicated by the negative correlation with PD and m_75_ and the related inverted association with breast cancer.

There has been considerable work in relating textures to breast cancer risk [9, 19]. We are unaware of techniques that are similar to this correlation approach. Metrics derived from the co-occurrence approach may be considered as distant relatives. When developing co-occurrence metrics [9, 20], spatial correlation is considered by selecting specific spatial distances and directions between two relative locations (i.e. some number of pixel distances in the x-direction and some number of pixel distances in the y-direction) resulting in metrics based on bivariate histograms for fixed distances. In contrast, our approach considered correlation as a function of all distances and directions up to the specific local region size. To our knowledge, the use of a template approach and zero padding to mitigate artifacts of the discrete FT in this context are novel. The template comparison is the important link connecting the local Fourier analysis (i.e. the correlation function) with the summary measurements; this essentially transforms a parametric approach to a non-parametric approach. Both correlation measures provided breast cancer associations similar in magnitude to those provided by PD. The longer spatial range m_75_ measure was moderately correlated with PD, whereas the s_25_ measure showed a weak correlation with PD.

Our study design has several limitations. The Study 1 exploration resulted in many measures. The method of selecting measures for evaluation in Study 2 may be less than optimal. This technique relied on univariate t-test findings, which are not the final metrics of interest (i.e. ORs). As such, selecting measures for further analysis could have some deficiencies. For example, image metrics can be influenced by BMI or other factors. We have noted that controlling for BMI either has a negligible impact or strengthens a given image measure’s breast cancer association. As illustration, we have provided a supplemental table where BMI and ethnicity were controlled for separately for m_75_ and s_25_. It interesting to note that m_75_ does not have a significant association in Study 2 until controlling for BMI in isolation. Sampling of cases and controls was not population-based, but rather a mixture of cases derived from an NCI-designated comprehensive cancer center inclusive of referrals from the community. The current findings should be replicated in population-based studies, although there is no evidence indicating the cases are not representative or referral to MCC is based on the images that formed the basis of the current work. Our analysis was restricted to areas that approximate the largest rectangle that fits within the breast region. Thus, a portion of the breast area was not included in the spatial correlation analysis.

The comparisons with PD suggest that this influence is likely to be negligible. The demographic of our population was primarily Non-Hispanic Caucasian women. Further studies with a wider range of race and ethnicity should be implemented to explore whether these measures generalize to wider populations for risk prediction purposes. As illustrated in Figure 4 (bottom row), within a given mammogram the correlation properties vary from region to region. Our study did not attempt to quantify this nonstationary statistical behavior beyond summarizing the local behavior to produce a given measure.

## 5. Conclusion

Mammograms contain content informative of the future risk of breast cancer, although the basis for this connection is not understood [15]. Most often, the degree of dense tissue is used for risk prediction purposes. Measurements that focus on other image attributes, such as those presented in this report, rather than breast density directly may be useful for informing future studies designed to unravel the biology of risk. A given correlation metric is defined over a specific spatial distance range, which provides another measurable parameter that can be used for analytical purposes. Whether metrics of this kind provide additional information in the mammographic-risk landscape will require more elaborate studies in the future. On the other hand, this correlation approach is a general analysis technique. By hypothesis, this approach may be useful for other forms of image analyses with varying endpoints beyond mammography, noting the template can be easily modified.

## Acknowledgements

This work was supported by the National Institutes of Health grants R01CA114491, R01CA166269, and U01CA200464.

