## Supplemental Table for "Spatial Correlation and Breast Cancer Risk"

| m_75_ (Study 1) | | | | | |  | s_25_ (Study 1) | | | | | |
| --- | --- | --- | --- | --- | --- | --- | --- | --- | --- | --- | --- | --- |
| LOG CON | LOG STD | unadjusted OR (95% CI) | ethnicity adjusted OR (95% CI) | BMI adjusted OR (95% CI) | BMI and ethnicity adjusted OR (95% CI) |  | LOG CON | LOG STD | unadjusted OR (95% CI) | ethnicity adjusted OR (95% CI) | BMI adjusted OR (95% CI) | BMI and ethnicity adjusted OR (95% CI) |
|  | 0.1774 | 0.85 (0.75, 0.96) | 0.85 (0.74, 0.96) | 0.70 (0.60, 0.81) | 0.69 (0.60, 0.81) |  |  | 0.0867 | 1.22 (1.07, 1.38) | 1.21 (1.06, 1.37) | 1.31 (1.15, 1.49) | 1.30 (1.14, 1.49) |
| Az |  | 0.55 (0.51, 0.59) | 0.60 (0.56, 0.63) | 0.60 (0.56, 0.64) | 0.64 (0.61, 0.68) |  | Az |  | 0.54 (0.50, 0.58) | 0.59 (0.55, 0.63) | 0.59 (0.55, 0.63) | 0.63 (0.59, 0.67) |
| m_75_ (m_37_ Study 2) | | | | | |  | s_25_ (s_13_ Study 2) | | | | | |
| LOG CON | LOG STD | unadjusted OR (95% CI) | ethnicity adjusted OR (95% CI) | BMI adjusted OR (95% CI) | BMI and ethnicity adjusted OR (95% CI) |  | LOG CON | LOG STD | unadjusted OR (95% CI) | ethnicity adjusted OR (95% CI) | BMI adjusted OR (95% CI) | BMI and ethnicity adjusted OR (95% CI) |
|  | 0.1333 | 0.80 (0.62, 1.02) | 0.80 (0.62, 1.02) | 0.69 (0.52, 0.90) | 0.67 (0.51, 0.89) |  |  | 0.0538 | 1.29 (1.03, 1.63) | 1.29 (1.03, 1.62) | 1.34 (1.06, 1.70) | 1.34 (1.06, 1.69) |
| Az |  | 0.55 (0.48, 0.62) | 0.55 (0.48, 0.62) | 0.62 (0.55, 0.69) | 0.61 (0.54, 0.68) |  | Az |  | 0.56 (0.48, 0.63) | 0.57 (0.49, 0.64) | 0.63 (0.56, 0.70) | 0.63 (0.56, 0.70) |

**Supplemental Table**: Expanded conditional logistic regression modeling for m_75_ and s_25_. This table provides odd ratios (ORs) and the area under the receiver operating characteristic curve (Az) with 95% confidence intervals (CIs) for the two selected spatial correlation metrics for both studies: m_75_ (or equivalently m_53_ for Study 2) and s_25_ (or equivalently s_17_ for Study 2). Models are provided in unadjusted, adjusted for ethnicity in isolation, adjusted for body mass index (BMI) in isolation, and adjusted for BMI and ethnicity in tandem formats. This table complements Table 4.
